# Intraneuronal binding of amyloid beta with reelin - implications for the onset of Alzheimer’s disease

**DOI:** 10.1101/2023.10.30.564686

**Authors:** Asgeir Kobro-Flatmoen, Stig W. Omholt

## Abstract

It was recently shown that in anteriolateral entorhinal cortex layer II neurons (ECLII neurons) in McGill-R-Thy1-APP homozygous transgenic rats (a model commonly used to study Alzheimer’s disease (AD)), the glycoprotein reelin and intracellular amyloid-*β* (A*β*) engage in a direct protein–protein interaction. Numerous studies of the human brain supported by experimental results from rodent and cell models point to a role for intracellular oligomeric A*β* in the onset of AD. If reelin functions as a sink for intracellular A*β* and if the binding to reelin makes A*β* physiologically inert, it implies that reelin may prevent the neuron from being exposed to the detrimental effects typically associated with oligomeric A*β*. Considering that reelin expression is extraordinarily high in the major subset of ECLII neurons compared to most other cortical neurons, such a protective role appears very difficult to reconcile with the fact that ECLII is clearly a major cradle for the onset of AD in humans. Here we show that this conundrum may be resolved if ECLII neurons have a much higher maximum production capacity of A*β* than neurons expressing low levels of reelin. We provide a rationale for why this difference has evolved, and argue that the higher maximum production capacity of A*β* in ECLII neurons may in a senescent A*β*-inducing physiology predispose these neurons to initiate AD development.

**Author summary:** Amyloid-*β* is a small peptide that is widely recognized as one of the main culprits involved in the development of Alzheimer’s disease. It was recently shown that in the major subset of neurons in entorhinal cortex layer II, which expresses high levels of the protein reelin, amyloid-*β* and reelin bind to each other. These neurons, which are strongly involved in memory formation, are among the first to die in subjects with Alzheimer’s disease. If intracellular amyloid-*β*, which is clearly involved in the onset of the disease, becomes physiologically inert when it binds to reelin, it implies that reelin can prevent the neuron from being exposed to the detrimental effects of increased levels of amyloid-*β*. Considering that reelin expression is extraordinarily high in ECLII neurons compared to most other cortical neurons, such a protective role appears very difficult to reconcile with the fact that ECLII constitute the predominant cortical site for initiation of Alzheimer’s disease. Here, we show that this paradox may be resolved if ECLII neurons have a much higher maximum amyloid-*β* production capacity than neurons expressing low levels of reelin. We provide reasons why this difference has evolved and argue that it, in a senescent physiology, predisposes ECLII neurons to initiate the development of Alzheimer’s disease.

## Introduction

Alzheimer’s disease (AD) is a neurodegenerative brain disease that causes dementia. Two proteins have been identified as central in the development of the disease, namely amyloid-*β* (A*β*) and tau. Studies of large human cohorts strongly suggest that the disease starts with an increase in A*β* in the brain, leading to hyperphosphorylated tau (p-tau) and eventually to the formation of insoluble p-tau aggregates, called neurofibrillary tangles (NFTs) [1, 2].

It is still somewhat underappreciated that the entorhinal cortex (EC), located adjacent to the hippocampus in the medial temporal lobe and being the main gateway of information flow between the neocortex and the hippocampus, is a major cradle for the onset of AD [1]. It plays a crucial role in the formation, consolidation, and retrieval of memories and spatiotemporal representation, and long before the clinically recognized symptoms of AD appear, such as substantial or persistent memory loss, it is already severely degenerated [3–5]. The cortical onset of NFTs is predominantly restricted to the lateral or anteriolateral portion of the EC (alEC), in projection neurons that reside superficially in layer II (ECLII) [2, 6, 7]. Since this population of neurons shows clear signs of degeneration from the earliest stages of AD, discovering the mechanisms responsible for this impairment is arguably of considerable importance if we want to achieve a truly efficient therapy against AD.

The function of the large glycoprotein reelin in adult mammalian brains includes synaptic modulation, induction of enhanced spine density, and promotion of long-term potentiation [8]. Most layer II projection neurons in alEC are unique among cortical excitatory neurons by expressing high levels of reelin (dubbed Re^+^ECLII neurons in the following). In terms of existing data and interpretations of AD development, at least three possibly interlinked paradoxes are attached to this observation.

The first paradox arises from the recent observation that in alEC layer II neurons in McGill-R-Thy1-APP homozygous transgenic rats (a model commonly used to study AD), reelin and intracellular A*β* engage in a direct protein–protein interaction [9]. Considering that numerous studies of the human brain using live imaging, immunohistochemistry, and biochemistry, supported by experimental results from rodent and cell models, point to a role for intracellular A*β* in non-fibrillated forms in the onset of Alzheimer’s disease [1], this is an intriguing discovery, as it suggests that reelin can function as a sink for intracellular A*β*. If we assume that the A*β*-reelin complex is physiologically inert, this would imply that ECLII neurons with a high reelin level would be protected from the detrimental effects associated with increased intracellular A*β* expression. This does not resonate with the observation that these neurons are among the first to die when the AD phenotype unfolds.

The second paradox arises from our current knowledge of reelin signaling and p-tau formation. Reelin is likely crucial for memory formation and, upon binding to its main receptor in the brain, ApoER2, it triggers a signaling cascade that enhances glutamatergic transmission [8]. Additionally, the binding of reelin to ApoER2 triggers a signaling cascade that strongly inhibits the activity of glycogen synthase kinase 3*β* (GSK3*β*) [10–12]. GSK3*β* is one of the main kinases that phosphorylates tau, and constitutive tau phosphorylation causes the formation of NFTs. It has been experimentally demonstrated that interaction with A*β* reduces the capacity of reelin to inhibit GSK3*β* kinase activity [13]. This suggests that a high intracellular reelin level would buffer against NFT formation in a senescent physiology that causes pathological expression of A*β*. But this does not resonate with the observation that Re^+^ECLII neurons are typically the first to form NFTs in Alzheimer’s disease.

The third paradox arises from the relationship between reelin function and the frequently advocated hypothesis that infectious agents contribute to the pathogenesis of AD. An infection basis for AD has been a contentious issue for decades. However, during the last decade, a substantial body of data has accumulated that at least supports that A*β* has antiviral [14, 15] and antimicrobial properties [16, 17] properties. And they support that A*β* oligomers are capable of binding to viruses [15, 18] as well as microbes [19]. Furthermore, the antimicrobial properties of A*β* have been reported to be reminiscent of the canonical human antimicrobial peptide LL-37, showing antimicrobial activity equivalent to or, in some cases, superior to this cathelicidin [20] against several clinically relevant bacteria [16]. Therefore, several researchers have proposed that A*β* participates in an immune response to acute microbial or viral infection of the brain [21–24]. Assuming that such an immune response would trigger an increase in the expression of A*β* during pathogen infection, it follows that in ECLII neurons with constitutively high reelin levels, A*β* would be prevented from exerting its immunological function, as it would become attached to reelin. Although the actual transport mechanisms and subsequent spread of viral infections through the central nervous system (CNS) are still poorly understood [25], the available data clearly suggest that ECLII neurons can be exposed to viral infection through their connection to olfactory receptor neurons and serve as a portal for further CNS infection [25–27]. Furthermore, the EC receives vascular input from the posterior and middle cerebral arteries, which forms particularly dense reticulated networks around individual clusters of ECLII neurons. Together with the neuron clusters themselves, these networks give rise to the characteristic bumps on the surface of the EC, known as the entorhinal verrucae. This redundancy in vascular input suggests that EC may also be particularly exposed to bloodborne pathogens [28]. Due to this propensity to become exposed to pathogens and assuming that increased A*β* production is indeed part of an intraneuronal immune response under native physiological conditions, it does not make evolutionary sense that reelin-expressing ECLII neurons should be more vulnerable to pathogen infection than other neurons.

Anticipating that resolving the third paradox may provide the key to resolving the other two, we constructed a simple mathematical model that describes the dynamics between intracellular A*β*, reelin and p-tau formation through the GSK3*β* pathway as a function of short-term pathogen exposure. We found that if the maximum possible production rate of A*β* is several times higher in reelin-expressing ECLII neurons than in other cortical neurons, Re^+^ECLII neurons are capable of invoking an immune response similar to that of other cortical neurons that do not produce high levels of reelin. We use the model to predict how Re^+^ECLII neurons that carry genotypes on opposite sides of the AD risk spectrum will respond to an elevated production rate of A*β*. And we use the immune response results to argue that in a senescent physiology, their higher A*β* production capacity and their very high energy requirement [1, 29] can render these neurons extraordinarily vulnerable to AD development.

## Materials and methods

### Description of the dynamic model

The model consists of four differential equations that describe the time rate of change of the amount of free intracellular A*β* [A*β*], the amount of free intracellular reelin [Reelin], the amount of intracellular A*β* bound to reelin [A*β*_*Reelin*_], and the amount of phosphorylated GSK3*β* that is inhibited from inducing hyperphosphorylated tau production (GSK3*β*_*p*_). The four differential equations are as follows.

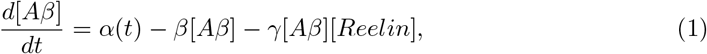

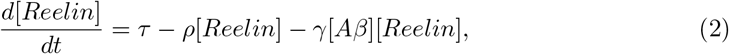

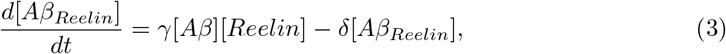

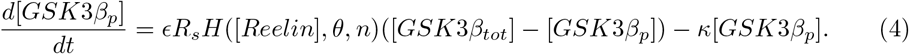

Equation (1) says that the time rate of change of free A*β* is a function of the production rate of A*β* at time *t* (*α*(t)), a first-order decay of A*β* (*β* = constant), and a second-order binding reaction between A*β* and reelin (*γ* = constant). We assume monomeric binding of reelin to A*β*. The decay term may also include extracellular transport.

Equation (2) says that the time rate of change of free reelin is a function of the production rate of reelin (assumed to be constant in a given neuron type), a first-order decay of free reelin (*ρ* = constant) and a second-order binding reaction between A*β* and reelin (identical to the last term in Eq. (1)). Due to the high intracellular concentration, it is very likely that reelin is produced in Re^+^ECLII neurons. However, we do not know whether this is the case in neurons with low constitutive levels of reelin. It may be that the reelin in these cells is delivered by interneurons [30]. In this case, the production term will have to be viewed as an import term. However, this does not affect the model as long as all low-reelin neurons are provided with about the same amount of reelin per time unit. We performed a semiquantitative analysis of in-house image data that showed that all neurons possess at least a reelin signal that is clearly above the background signal (data not shown).

Equation (3) says that the time rate of change of the amount of the A*β*_*Reelin*_ complex is a function of its production rate (identical to the last term in Eq. (1)) and first-order decay of the complex (*δ* = constant). The decay term may also include extracellular transport. We do not account for the possibility that A*β*_*Reelin*_ complexes aggregate into configurations that may make them less exposed to intracellular degradation or export, warranting a more complex decay/export term.

Equation (4) says that the time rate of change of the amount of GSK3*β* prevented from inducing tau hyperphosphorylation is given by the rate of GSK3*β* phosphorylation due to reelin signaling times the amount of unphosphorylated GSK3*β* minus the rate by which GSK3*β* loses this inhibition (which is proportional to the amount of inhibited GSK3*β* (*κ* = constant)). The design of the expression describing the rate of phosphorylation is motivated by the fact that reelin inactivates GSK3*β* by binding to ApoER2 as a dimer [31]. Binding induces a receptor cycling process that activates a signaling cascade that eventually causes phosphorylation of GSK3*β* [25], inhibiting its kinase activity. We assume that above a certain level of free reelin, the receptor cycling rate saturates, i.e., practically speaking, above this level there is almost no additional effect of reelin on the GSK3*β* phosphorylation rate and therefore the rate becomes almost constant. It is reason to believe that the functional form of the saturation process is more sigmoidal than hyperbolic because reelin binds to ApoER2 as a dimer and because chemical reactions where one of the reactants is bound to a surface cause so-called fractal kinetics where even reactions involving no cooperativity will behave as they did [32–34]. To capture such a relationship, the Hill function 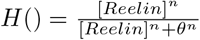 is very useful. Here, the steepness of the sigmoidal function is regulated by the parameter *n* and the threshold value of reelin where it engages 50% of the ApoER2 population in inhibitory signaling is regulated by the parameter *θ*. The constant *R*_*s*_ is the amount of free reelin where the Hill function value is greater than 0.99, i.e., very close to saturation. Thus, the product *R*_*s*_ x H() can be viewed as a function that describes the effective amount of intracellular free reelin in terms of signaling strength. Since there is no further phosphorylation when all GSK3*β* molecules are phosphorylated, the term ([*GSK*3*β*_*tot*_] *−* [*GSK*3*β*_*p*_] captures that even when reelin signaling is high, the phosphorylation rate will decrease with decreasing amount of unphosphorylated GSK3*β*. The parameter *ϵ* is constant and [*GSK*3*β*_*tot*_] is the total intracellular amount of GSK3*β*, assumed to be constant. The way the GSK3*β* phosphorylation rate in Eq. (4) varies with the reelin level and the proportion of phosphorylated GSK3*β* is depicted in Fig. 1 as a normalized 2-d heat map.

**Fig 1.**
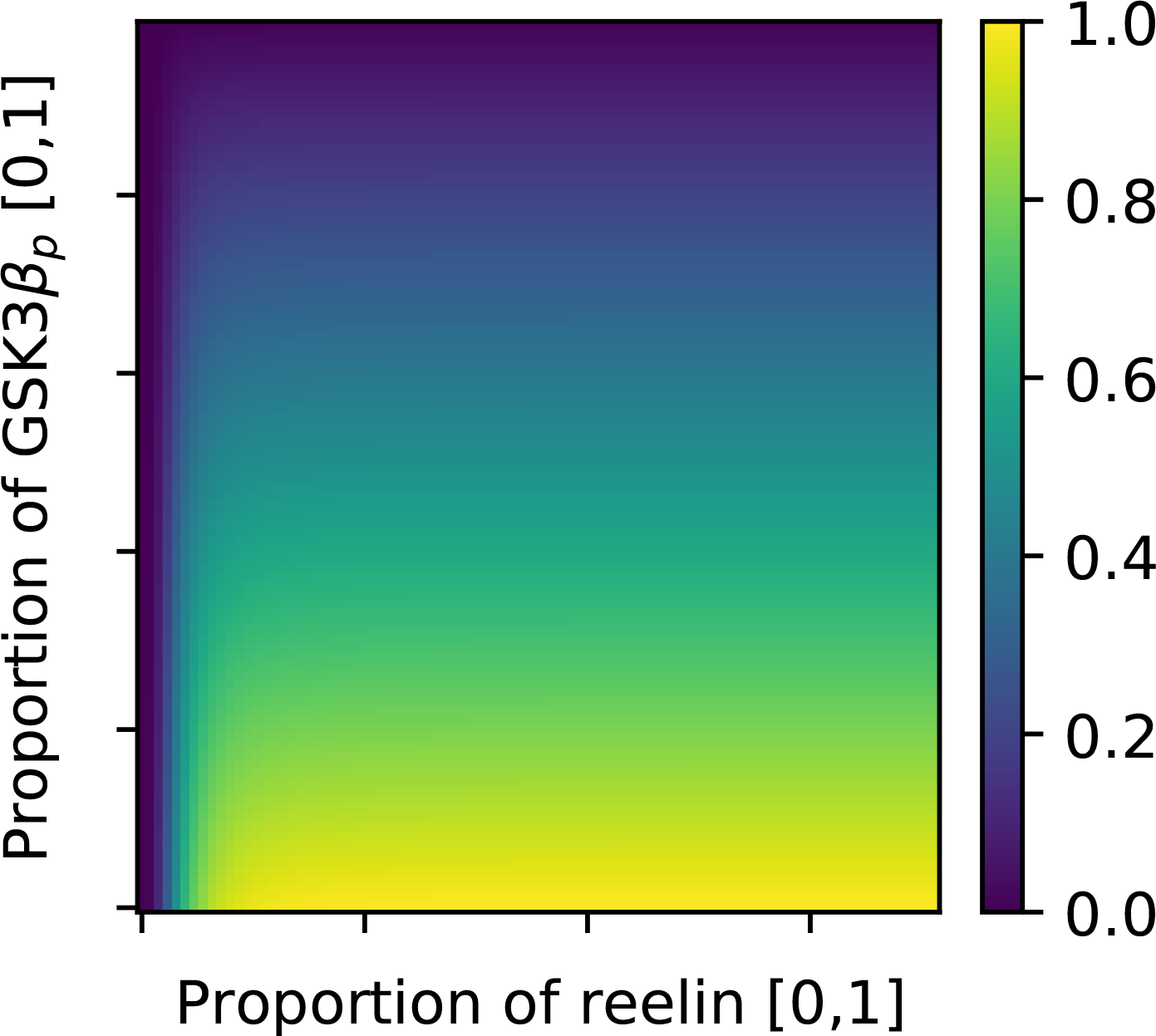
GSK3*β* phosphorylation rate. The figure shows how the term describing the GSK3*β* phosphorylation rate in Eq. (4) varies with the cellular reelin level (normalized) and the proportion of already phosphorylated GSK3*β* molecules. See the text for further explanation.

We assume that Eqs. (1)-(4) capture the dynamics of the four state variables to a reasonable degree in Re^+^ECLII neurons and in cortical neurons that possess low constitutive levels of reelin (called typical cortical neurons in the following). It implicitly assumes that the latter group of neurons produces (or receives) sufficient amounts of reelin to suppress the activation of p-tau formation through GSK3*β* under native physiological conditions and that this suppression begins to fail when the intracellular level of free reelin falls below a certain threshold. Furthermore, we assume that the two types of neurons have identical GSK3*β* regulation. That is, the kinetic parameters underlying the inhibition of GSK3*β* kinase activity are the same in terms of the amounts of ApoER2, GSK3*β* (and the signaling molecules located between these two), and in terms of the functional relationships that determine the propagation of the signal between reelin and GSK3*β*. Similarly, we assume that the binding affinity between reelin and A*β* and the decay constants of A*β*, A*β*_*Reelin*_, reelin, and GSK3*β*_*p*_ are the same in both groups of cells. The only two parameters in the model that are not assumed to be identical between the two groups are the reelin production rate (to account for the observed difference in the reelin level) and the production rate of A*β* during infection (explained below).

### Initial parameterization of the model

Due to the lack of data, we had to assume intracellular half-life times for free A*β*, free reelin, A*β*_*Reelin*_ and GSK3*β*_*p*_. Given the purpose of the model, it is sufficient with quite approximate estimates. We assumed the half-life of free A*β* and free reelin to be 6.9 hours, i.e., *β* = *ρ* = 0.1 *h*^*−*1^. This is about 75% of the median half-life of intracellular proteins in mammalian cells [35]. The intracellular half-life of A*β*_*Reelin*_ is likely to be substantially higher and we set it to be about 69 hours, i.e., *δ* = 0.01 *h*^*−*1^. We let *γ* = 6, and preliminary simulations of the model showed that the results were quite insensitive to the value of this parameter describing the binding affinity between reelin and A*β*. We assumed that the intracellular amount of GSK3*β* is under homeostatic control, and that the half-life of the phosphorylated state of GSK3*β* due to reelin signaling is approximately 1.5 hours, giving *κ* = 0.45 *h*^*−*1^. The Hill function parameter *n* in Eq. (4) was set equal to 3 in accordance with our above justification of Eq. (4). The values of *θ* and *R*_*s*_ were calibrated according to the expression span of reelin (*θ* = 4 and *R*_*s*_ = 16 in the simulations underlying Figs. 1, 2 and 3).

**Fig 2.**
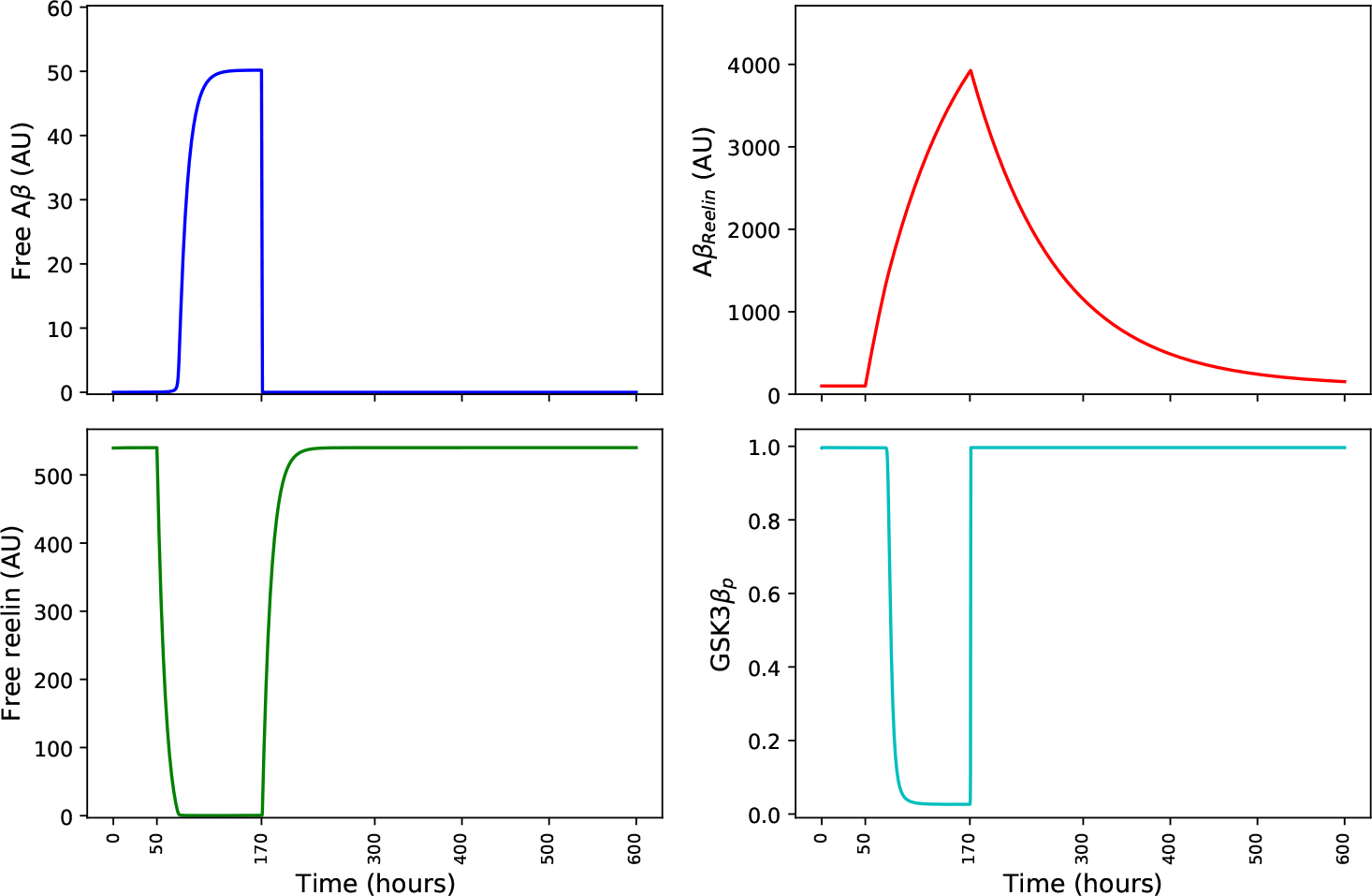
Immune response in Re^+^ECLII neurons. The four panels show how the levels of free A*β*, A*β*_*Reelin*_ and free reelin and the proportion of phosphorylated GSK3*β* develop during an infection event that lasts 120 hours (from t=50 to t=170). AU = artificial units. Parameter values: *α*=1 if t*≤*50 or t*>*120, else *α*=60; *β*=0.1; *γ*=6; *δ*=0.01; *τ* =55; *ρ* = 0.1; *ϵ*=8; *n*=3; *θ*=4; *G*_*tot*_=100; *R*_*s*_=16; *κ*=0.45. See the text for further explanation.

**Fig 3.**
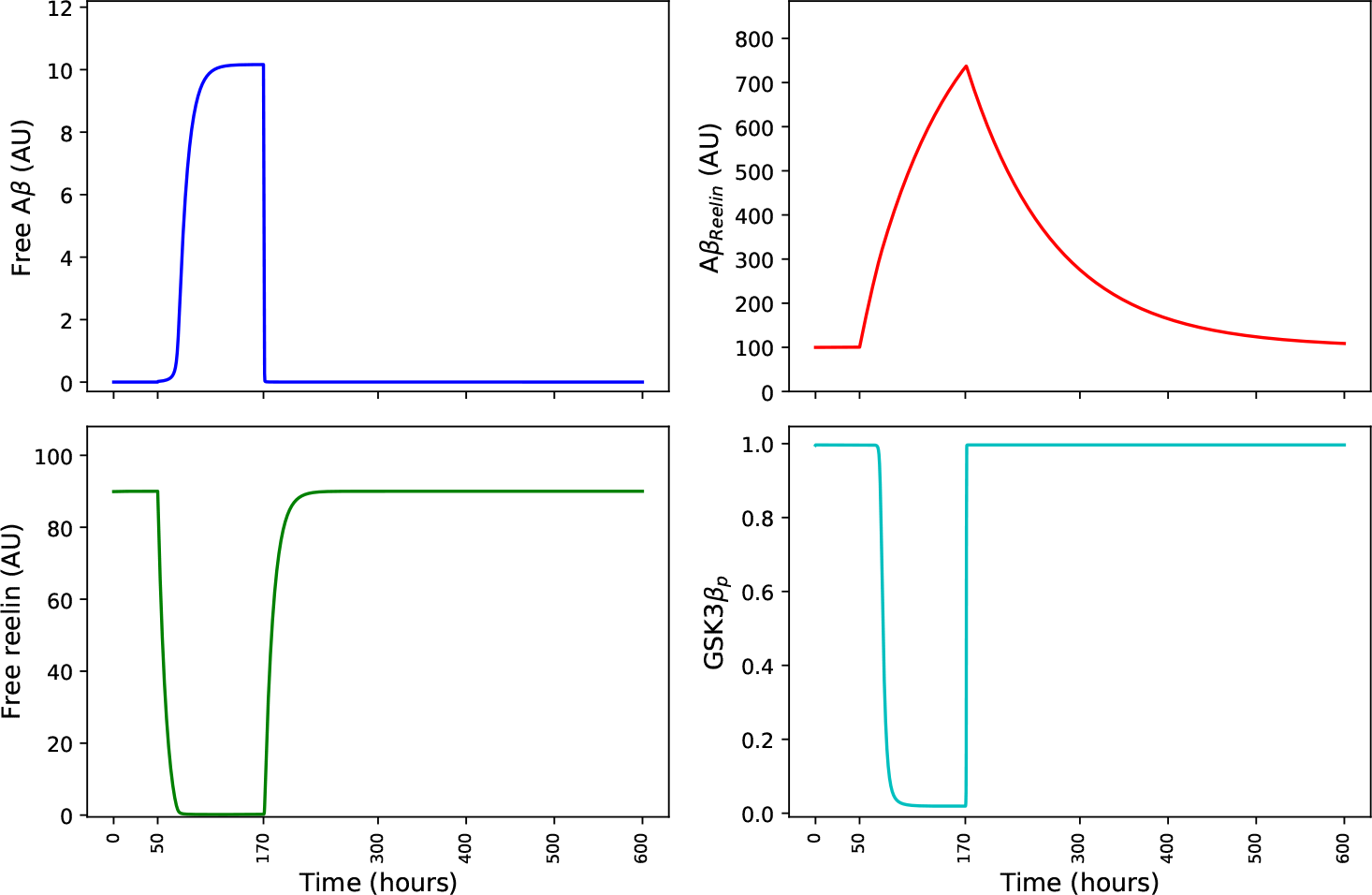
Immune response in typical cortical neurons. The four panels show how the levels of free A*β*, A*β*_*Reelin*_ and free reelin and the proportion of phosphorylated GSK3*β* develop during an infection event that lasts 120 hours (from t=50 to t=170). Parameter values: *α*=1 if t*≤*50 or t*>*120, else *α*=11; *β*=0.1; *γ*=6; *δ*=0.01; *τ* =10; *ρ* = 0.1; *ϵ*=8; *n*=3; *θ*=4; *G*_*tot*_=100; *R*_*s*_=16; *κ*=0.45. Phosphorylated GSK3*β* is normalized against *G*_*tot*_. See the text for further explanation.

### The feedback structure of the model

Equations (1)-(4) can be solved analytically to give the steady state values when *α*(t) = constant. We are not showing these expressions here as they are not very informative, but they were used to find the steady state values in the baseline condition with a low and constant *α* value. The time evolution of the system (see below) could have been approximated by using the steady state solutions for a low and a high *α* value, but we have not pursued this. The Jacobian 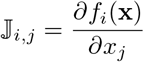 of the system is:

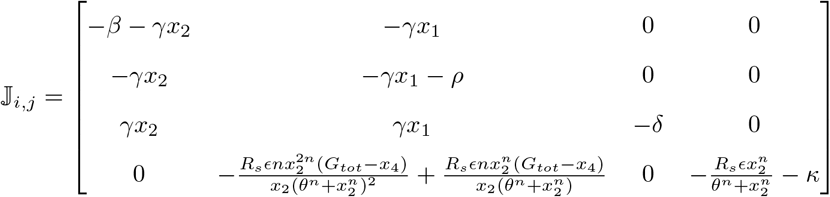

We see from the Jacobian [36] that the system contains four negative autoregulatory loops (one for each state variable) and one positive feedback loop between *Aβ* and reelin (i.e., the sign of element (1,2) x element (2,1) is positive). We have found it challenging to further simplify the model without losing essential features.

The model was coded in Python in a Jupyter lab environment, and SymPy, a Python library for symbolic mathematics, was used to calculate closed-form expressions for the steady states of the four state variables under native physiological conditions. The Python library SciPy was used to solve Eqs. (1)-(4) numerically.

## Results and Discussion

### Mimicking an immune response in the two groups of neurons

If A*β* is an important player in an intraneuronal immune response against pathogen infection under native physiological conditions, and based on what we now know about the relationship between reelin and A*β* and the relationship between reelin and p-tau formation, we believe that the following scenario is likely in all cortical neurons: Production of A*β* is very low as long as the cell does not detect any infection and there will be a marginal accumulation of A*β*_*Reelin*_ complexes. An infection will induce an abrupt increase in production of A*β*. This will lead to increased accumulation of A*β*_*Reelin*_, depletion of the free reelin level, increase of the free A*β* level, and marked p-tau formation. After eliminating the pathogen, the cell reduces its A*β* level to its low baseline level, which eventually leads to the restoration of the original levels of free reelin and A*β*_*Reelin*_ and to the prevention of further p-tau formation through the reelin - GSK3*β* pathway. We then set out to determine whether the dynamic model was capable of recapitulating this scenario in cortical neurons with, respectively, high or low levels of constitutive reelin expression.

We initially let a neuron produce A*β* (*α*(*t*)) at a constant low rate. The initial values of the four state variables at time *t*_0_ were set identical to the steady state values under the given parameter regime. These values were found analytically by solving Eqs. (1-4) after setting the left-hand sides to zero. At an arbitrary time *t*_1_ (50 hours), we assumed that the cell detected a pathogen invasion and immediately increased its A*β* production rate to a constant level several times higher. Except for presuming its existence, we were agnostic about the details of the sensing mechanism and how it induces an increase in A*β* production. At time *t*_2_ (170 hours), we assumed that the cell detected that it had eliminated the pathogen and that the production rate of A*β* immediately fell back to the low basal level. Again, we were agnostic about the details of this sensing mechanism and how it induces deactivation of A*β* production. We assumed that the infection lasted five days. However, this figure is not important, as shorter or longer duration will only affect the size of the accumulated A*β*_*Reelin*_ pool and therefore the time it will take before the pool size returns to normal levels after the cessation of infection. Note that we deliberately assume that reelin production is not turned off or down-regulated during an infection event. Although the available data do not seem to support such regulation, we cannot exclude this possibility, and it would in case have a strong impact on our conclusions.

As long as we are only interested in identifying the prerequisites for achieving a sensible and similar immune response in both types of neurons under the assumption that the basic regulatory architecture is the same, we decided that this very simple way of describing a hypothetical immune response was fit for purpose.

In the case of Re^+^ECLII neurons with a high steady-state reelin level, without any loss of generality, we fixed the constitutive production rate of reelin (*τ* = 55). To mimic the initiation of the immune response after 50 hours, we increased the A*β* production rate (keeping all other parameters fixed) until it caused a drop in the level of free reelin large enough to induce at least a 97% reduction in the GSK3*β* phosphorylation degree. By this procedure, the model produced an immune response pattern fully in line with the scenario described above (Fig. 2). Note that under the given parameter regime, the intracellular amount of A*β*_*Reelin*_ does not reach steady state (Fig. 2) if the infection event lasts only five days.

We then assumed that the baseline free reelin level in typical cortical neurons that contain a low steady-state amount of reelin is approximately 17% of the level in Re^+^ECLII neurons and adjusted the reelin production rate (*τ*) accordingly. This figure is based on previous experience with immunocytochemical reelin staining [9, 37], indicating that a strong reelin signal reflects a reelin concentration that is at least 5-6 times higher than what is reflected by a signal that is slightly above background.

Except for the A*β* production rate, we kept all other parameters fixed. Similarly to Re^+^ECLII neurons, the rate of A*β* production during infection was adjusted so that it caused a drop in the level of free reelin large enough to induce at least a 97% reduction in the GSK3*β* phosphorylation degree. In this way, we also obtained a sensible immune response in this type of neuron (Fig. 3). We see that the free level of A*β* during infection is lower than in the Re^+^ECLII neurons. If we want the levels to be similar for reasons of immune response efficiency, this can be achieved, without notable changes in the other three state variables, by letting the A*β* production rate increase approximately 36%. In both cases, the immune response is very similar (Fig. 3) to that of Re^+^ECLII neurons (Fig. 2) in terms of the rapid decrease in reelin level and the degree of phosphorylation of GSK3*β* after the onset of infection.

In healthy Wistar rats, a low level of intracellular A*β*42 is found in neurons throughout the brain at all ages. The highest levels are found in ECLII neurons, in particular in neurons located in the superficial part of this layer, which is the part that contains the Re^+^ECLII neurons, and in hippocampal neurons at the CA1/subiculum border [38]. This suggests that the baseline production rate of A*β* is higher in Re^+^ECLII neurons than in almost all other cortical neurons. For simplicity reasons, we did not account for this when making Figs. 2 and 3, since equipping Re^+^ECLII neurons with a higher baseline production rate has a minor impact on the results. We do not know the reason for this elevated baseline production rate. It may be because more A*β* is needed for the normal operation of Re^+^ECLII neurons, or it may reflect that the configuration of the regulatory elements determining the A*β* production rate in Re^+^ECLII neurons is actually different from typical cortical neurons, leading not only to a higher A*β* production during infection, but also under baseline conditions.

In both groups of neurons, we deliberately tuned the production rate of A*β* during infection to be just above the threshold needed for the envisaged immune response to function. The effect of this tuning process on the four state variables can be visualized by varying the production rate of A*β* during infection between *±*50% relative to the original value, while keeping all other parameters fixed (Fig. 4, Re^+^ECLII neurons only).

**Fig 4.**
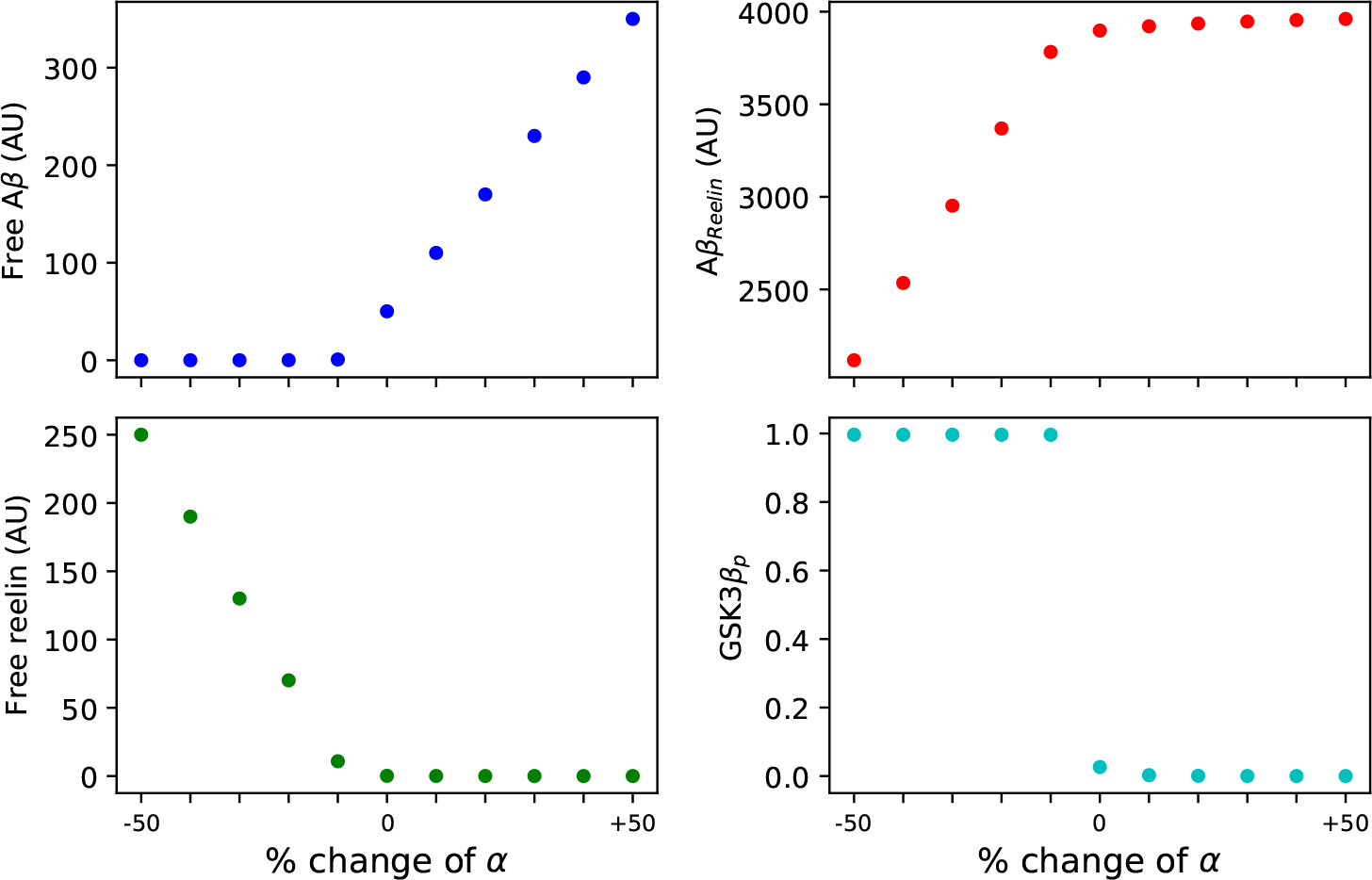
Sensitivity to change in A*β* production rate in Re^+^ECLII neurons. The four panels show how the levels of free A*β*, A*β*_*Reelin*_ and free reelin and the proportion of phosphorylated GSK3*β* during the infection period as a function of the value of *α* during infection. We let *α* vary ±50% relative to the original value, while keeping all other parameters fixed. Except for the actual values of the y axes in the first three panels, the graphs for typical cortical neurons are almost identical.

If cortical neurons possess an intraneuronal immune response against pathogens, the model suggests that a prerequisite for Re^+^ECLII neurons to produce an immune response on par with cells that have much lower constitutive levels of reelin is that they have a production rate of A*β* that is several times higher during infection. For both groups of neurons, the model predicts that during infection there will be a dramatic drop in the level of free reelin and that shortly after the end of infection, the amount of A*β*_*Reelin*_ will be several times higher than before infection (Figs. 2 and 3). It also predicts that shortly after cessation of infection, the amount of A*β*_*Reelin*_ will be several times higher in Re^+^ECLII neurons compared to typical cortical neurons (Figs. 2 and 3). We believe that all predictions can be tested by infecting wild-type rodents with, for example, herpes simplex virus 1 (HSV1) [25] and measure free reelin, free A*β* and A*β*_*Reelin*_ levels before, during and after infection by existing immunocytochemistry protocols [9]. If the amount of free A*β* during infection is much higher in Re^+^ECLII neurons than in typical cortical neurons, this will indicate that the higher level is just a consequence of the high production rate of A*β* needed to overcome the buffering effect of reelin.

### The effect of genotype on the immune response in Re^+^ECLII neurons

The propensity to develop sporadic AD is at least 8x higher in ApoE*ϵ*4/*ϵ*4 genotypes than in the most common ApoE*ϵ*3/*ϵ*3 genotypes [39], making the ApoE*ϵ*4/*ϵ*4 genotype the most pronounced genetic risk factor for acquiring sporadic AD. The causal basis for the observed difference between ApoE*ϵ*4/*ϵ*4 and ApoE*ϵ*3/*ϵ*3 subjects has become a much debated issue, and there are numerous explanations in the literature [40]. On the opposite side of ApoE*ϵ*4/*ϵ*4 in the AD risk spectrum, we have the recently discovered ‘COLBOS’ genotype, which refers to a rare mutation in the gene encoding reelin that has been shown to cause extreme resilience to AD even when the carrier also possesses the autosomal dominant familial AD (FAD) PSEN1-E280A mutation [41]. In both cases, it is pertinent to ask whether major causative effects of these alleles are located within the neurons themselves. Conditional on the availability of human neuronal cell cultures in which neurons express a phenotype very similar to Re^+^ECLII neurons [42], the model can be used to predict the outcome of a cell culture experiment in which cells carrying ApoE*ϵ*4/*ϵ*4, ApoE*ϵ*3/*ϵ*3 and COLBOS genotypes are exposed to a substantially enhanced intracellular level of A*β*.

The reelin mutation in the COLBOS case (in the following called the COLBOS genotype) clearly helps EC neurons maintain a reelin level that is sufficient to prevent the formation of p-tau, despite being exposed to a high A*β* load [41]. One possible explanation is that the mutation causes a dramatic reduction in the affinity between A*β* and reelin. Thus, the model can be used to predict the effect of this mutation by letting the value of the parameter *γ* become much less than that used to produce Figs. 2 and 3, while keeping all other parameters fixed and not including the effect of the PSEN1-E280A mutation.

Recapitulating the effect of the ApoE*ϵ*4/*ϵ*4 genotype is less straightforward. Reelin signaling involves endocytosis of ApoER2 receptors. Apolipoprotein E (ApoE) isoforms differentially affect the recycling of these endosomes. ApoE*ϵ*3-containing endosomes readily recycle back to the surface, while those containing ApoE*ϵ*4 remain trapped in vesicles [43]. Here, we assume that this vesicle trapping causes some reelin and ApoER2 to be taken out of circulation at a constant rate. We mimicked this process by adding a constant term *−ω* to the right-hand side of Eqn. (2). We did not modify Eqn. (4) to capture the trapping of ApoER2 as it could be accounted for by increasing *ω*.

In the in silico representation of the cell culture experiment with Re^+^ECLII neurons containing the three genotypes, we let the cells remain at baseline steady-state levels for 50 hours before increasing the intracellular level of A*β*. Existing data imply that the addition of A*β* in the picomolar range to the medium of neuron-like cells [44] is sufficient to induce increased A*β* production, suggesting the existence of a positive autoregulatory feedback loop. Given the existence of this feedback loop the A*β* production rate will soon saturate at its maximum attainable level after adding synthetic A*β* to the medium. In the numerical experiment, we therefore changed the A*β* production rate from its base level to its maximum level after 50 hours, and let the model run for 550 hours.

Two major predictions of the model are that cells with the COLBOS genotype will show no signs of p-tau formation after 550 hours with increased A*β* production and that their free reelin level will be higher than in the two other genotypes (Fig. 5). A third prediction is that the free A*β* level in the ApoE*ϵ*3/*ϵ*3 genotype will be much lower than the levels in the COLBOS and ApoE*ϵ*4/*ϵ*4 genotypes.

**Fig 5.**
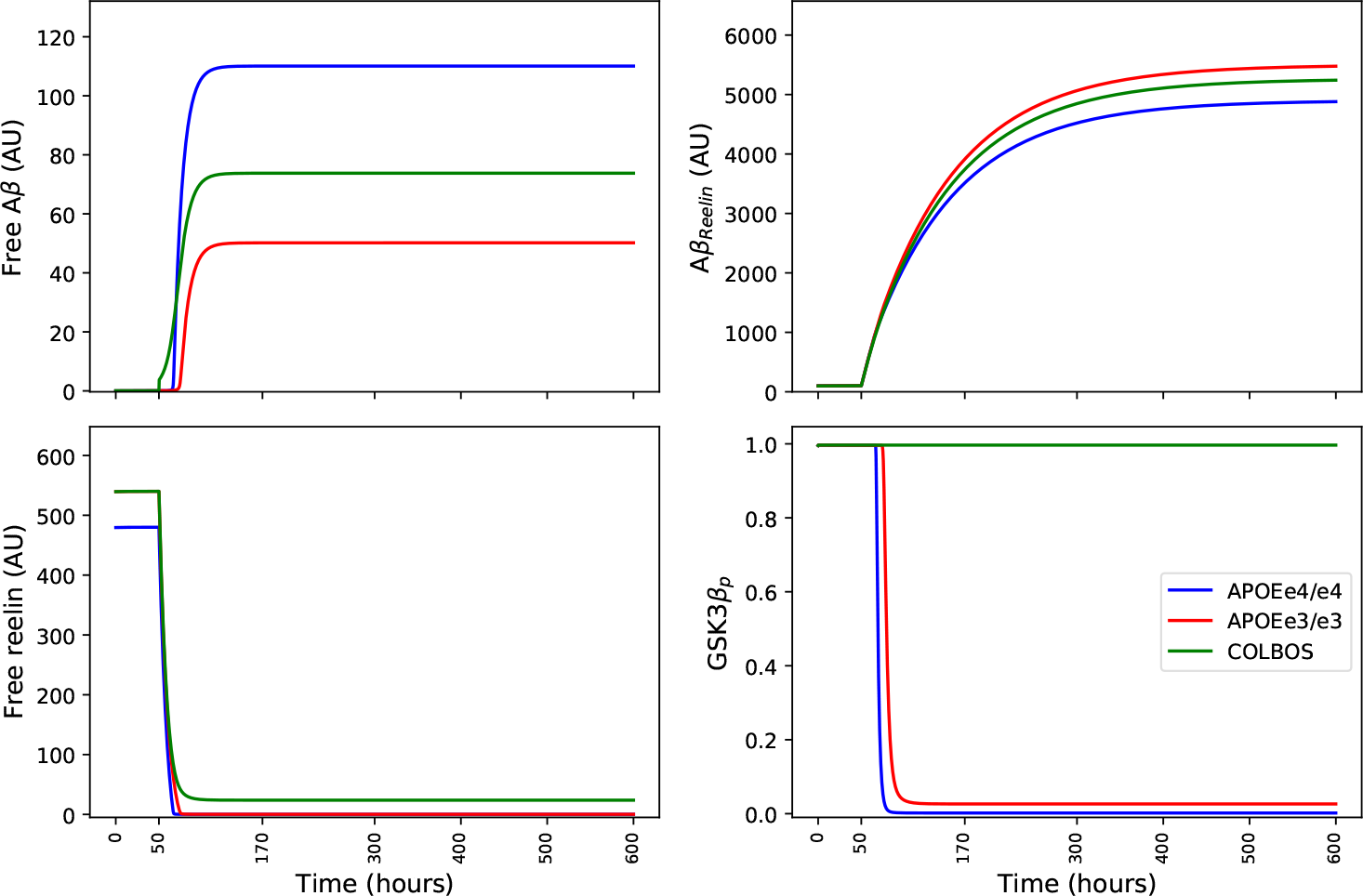
In silico cell culture experiment with Re^+^ECLII neurons having different genotypes. The four panels show how the levels of free A*β*, A*β*_*Reelin*_ and free reelin and the proportion of phosphorylated GSK3*β* are predicted to develop if one induces the neurons to produce high levels of A*β* from t=50 to t=600. The parameter values for the ApoE*ϵ*3/*ϵ*3 genotype were identical with those listed in the caption of Fig. 2. The parameter values for the COLBOS reelin mutation case are identical with those of the ApoE*ϵ*3/*ϵ*3 genotype, except that *γ*=0.03, i.e., we assume that the reelin mutation is genetically dominant. The parameter values for the ApoE*ϵ*4/*ϵ*4 genotype are identical with those of the ApoE*ϵ*3/*ϵ*3 genotype, but in this case we modified Eq. (2) by adding a constant term − *ω* = −6 to the right-hand side of the equation to account for the effect of vesicle trapping. See the text for further explanation.

The model robustly predicts that the ApoE*ϵ*4/*ϵ*4 and COLBOS genotypes will be exposed to much more A*β* than the ApoE*ϵ*3/*ϵ*3 genotype. In an in vivo context, this implies that more A*β* will be exported to the extracellular environment. Since both the COLBOS and ApoE*ϵ*4/*ϵ*4 genotypes are characterized by an extremely elevated amyloid plaque burden [41, 45], it appears that the model is capable of recapitulating this phenotype.

The in silico cell culture experiment supports the notion that the COLBOS genotype prevents p-tau formation by reducing the affinity between reelin and A*β*, which causes the free reelin level to stay above the threshold necessary for inhibiting GSK3*β* kinase activity even when the neuron is exposed to a severe free A*β* burden [41].

The results suggest that the formation of p-tau will be slightly higher in the ApoE*ϵ*4/*ϵ*4 genotype than in the ApoE*ϵ*3/*ϵ*3 genotype (Fig. 5). The difference is likely too small to have any functional significance. However, in a senescent physiology characterized by frequent bursts of increased A*β* production due to systemic inflammation or other causes, our results (Fig. 5) suggest that ApoE*ϵ*4/*ϵ*4 neurons are prone to be exposed to a much higher level of free A*β*, which implies that neurotoxic A*β* oligomers are far more likely to form. It remains to be determined how important this effect is compared to other functional effects of the ApoE*ϵ*4/*ϵ*4 genotype. However, it suggests that the vesicle trapping caused by this genotype may be of considerable importance.

### Why are Re^+^ECLII neurons a cradle for AD?

The above results do not resolve the first two paradoxes presented in the Introduction, i.e., they do not explain why Re^+^ECLII neurons are predominantly the first to show NFT formation and among the first to die. They suggest that in a senescent physiology predisposing to frequent inflammation-driven A*β* production bursts, Re^+^ECLII neurons will export much more A*β* into the extracellular environment than most other cortical neurons. However, since A*β* plaques do not typically appear in ECLII together with the fact that the COLBOS case shows that A*β* plaque burden can be extraordinarily high without invoking p-tau formation in ECLII [41], we suspect that an elevated export rate of A*β* in Re^+^ECLII neurons is not determinative.

The ROS-scavenging system is likely to have the same bandwidth in Re^+^ECLII neurons and in other cortical neurons. In native physiology, it is reasonable to assume that both experience the same ROS stress due to homeostatic regulation. However, due to their very high metabolic rate [1, 29], Re^+^ECLII neurons produce comparatively larger amounts of ROS through electron leakage from the electron transport chain (ETC) and from dysfunctional mitochondria before these are removed by mitophagy. Hence, Re^+^ECLII neurons are likely to have a more pronounced antioxidant defense signature than most other cortical neurons. This suggests that in the native condition Re^+^ECLII neurons are exposed to ROS stress that is comparatively closer to the threshold above which the ROS scavenging system would no longer be able to keep the ROS level within normal limits. When exposed to the same amount of supraphysiological oxidative stress, the actual intracellular ROS stress can therefore become disproportionally higher in Re^+^ECLII neurons than in typical cortical neurons. Since increased ROS stress is likely to induce an increase in A*β* formation, which in turn causes an additional increase in ROS stress [46], it follows from Figs. 2 and 3 that in a senescent physiology that causes frequent inflammation-driven A*β* production bursts, Re^+^ ECLII neurons can be exposed to much higher ROS stress (as well as other neurotoxic effects associated with A*β* oligomers). Since supraphysiological ROS levels can induce tau hyperphosphorylation [47] independently of the reelin-GSK3*β* pathway, it follows that the rate of NFT formation may become disproportionally higher in Re^+^ECLII neurons than in typical cortical neurons when exposed to an identical ROS-inducing inflammatory event (Fig. 6). Thus, recurrent senescence-driven oxidative stress appears to be capable of causing NFT accumulation earlier in Re^+^ECLII neurons than in typical cortical neurons.

**Fig 6.**
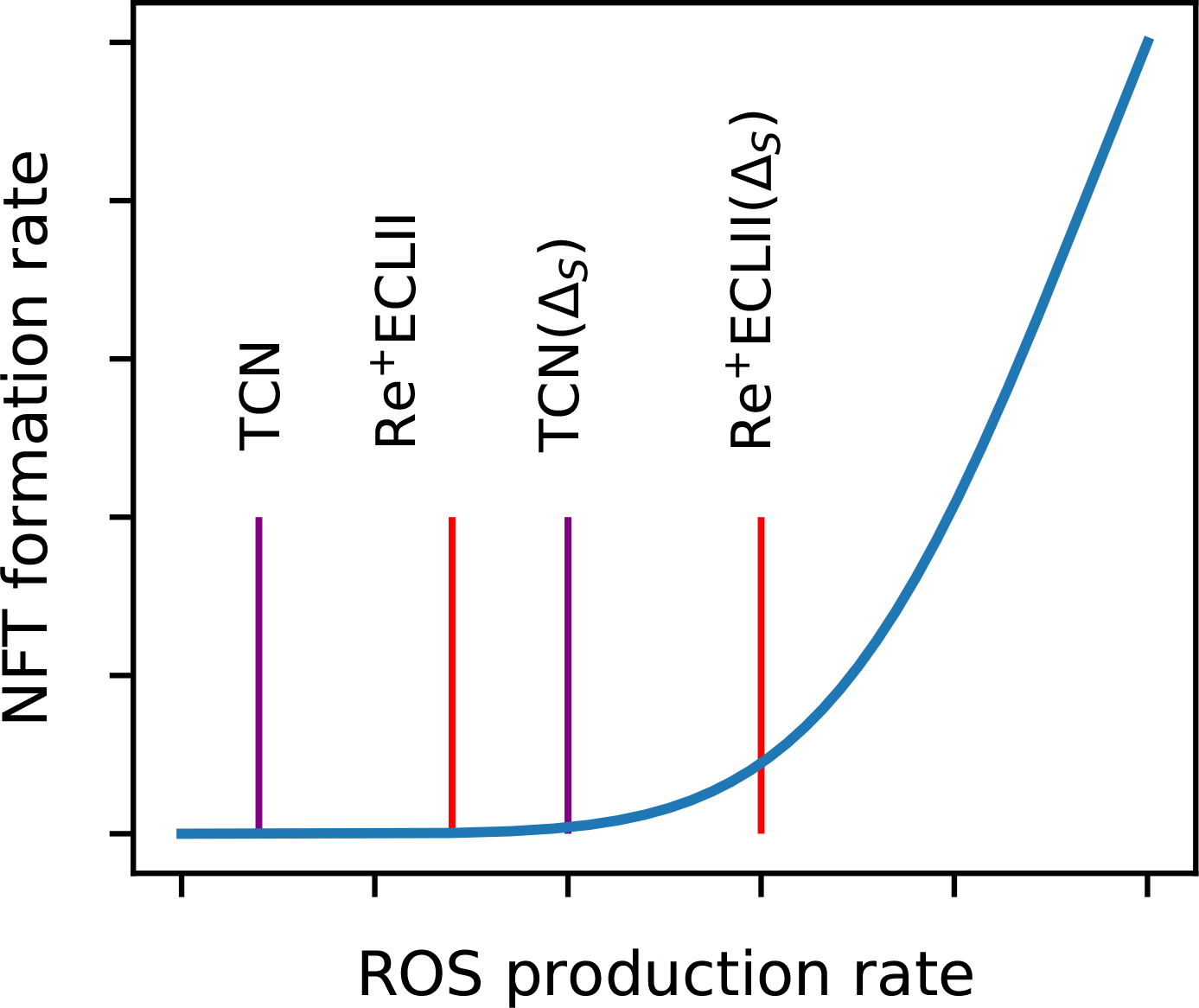
Possible relationships between ROS production rate and NFT formation rate in Re^+^ECLII neurons and typical cortical neurons (TCN) in a native physiology and in a senescent physiology causing elevated oxidative stress (Δ_*S*_). See the text for further explanation.

The above explanatory narrative is by no means likely to tell the whole story. But since it is completely consistent with the observation that Re^+^ECLII neurons are predominantly the first to show NFT formation and typically the first to die, it putatively resolves the two first paradoxes presented in the Introduction. And it supports our anticipation stated in the Introduction that resolving the third paradox would provide the key to resolving the other two.

### Concluding remarks

Although it has been established that A*β* has antiviral and antimicrobial properties, it remains to be conclusively shown that A*β* is indeed a major player in a concerted intraneuronal immune response. But if there is such a response, we claim that Re^+^ECLII neurons will have to sport a much higher maximum production capacity for A*β* than typical cortical neurons due to their extraordinarily high reelin production.

And we claim that this elevated production capacity is likely to be a causative factor in the very early etiology of AD. To support this claim, we have suggested mechanisms that may be sufficient to explain why Re^+^ECLII neurons are particularly vulnerable to the pathology that characterizes the development of sporadic AD despite the fact that their constitutive reelin level is several times higher than in typical cortical neurons.

In the model, we assumed that the decay of A*β*_*Reelin*_ is a first-order process. This may not be the case if A*β*_*Reelin*_ aggregates in such a way that if the amount of A*β*_*Reelin*_ becomes very large, the intraneuronal decomposition of A*β*_*Reelin*_ and the export of A*β*_*Reelin*_ to the extracellular environment stop following first-order kinetics due to a decrease in surface-to-volume ratio. In a senescent physiology characterized by frequent inflammatory events, this could cause Re^+^ECLII neurons to accumulate A*β*_*Reelin*_ far beyond the level they are designed to handle, leading to oxidative stress that causes both A*β* production [46] and tau hyperphosphorylation [47] independently of the reelin-GSK3*β* pathway. These responses would probably be sufficient to resolve the first two paradoxes alluded to in the Introduction. However, modeling such a causal chain of events is beyond the scope of this paper.

Neither have we considered that Re^+^ECLII neurons are clearly more prone to accumulate dysfunctional mitochondria very early in disease development than typical cortical neurons [48, 49]. We discuss this issue in more detail elsewhere, but note that this is likely related to their putatively extraordinarily high metabolic rate [1, 29] and that neurons with very high energy requirements are necessarily more prone to dysfunction when they experience a substantial drop in their energy supply. This implies the existence of additional cascading processes involving mitochondria that further drive EC to become a cradle for AD. The inclusion of such processes will require a much more comprehensive modeling effort, but we anticipate that the key premises underlying our model will continue to be valid.

## Acknowledgments

The authors wish to thank Eivind Almaas for his comments on a previous version of this manuscript. This work was supported by the K. G. Jebsen Foundation and the Kavli Institute for Systems Neuroscience Centre of Excellence Grant.

